# Persistence of Resident and Transplanted Genotypes of the Undomesticated Yeast, *Saccharomyces paradoxus* in Forest Soil

**DOI:** 10.1101/305060

**Authors:** James B. Anderson, Dahlia Kasimer, Clara Xia, Nicolas C. H. Shröeder, Patrick Cichowicz, Silvio Lioniello, Rudrackshi Chakrabarti, Eashwar Mohan, Linda M. Kohn

**Affiliations:** Department of Biology, 3359 Mississauga Road, University of Toronto, Mississauga Ontario L5L 1C6 Canada

**Keywords:** fungus, population, dispersal, genetic drift, 5FOA resistance

## Abstract

One might expect yeasts in soil to be highly dispersed via water or insects, forming ephemeral, genetically heterogeneous populations subject to competition and environmental stochasticity. Here, we report years-long persistence of genotypes of the yeast *Sacchaormyces paradoxus* in space and time. Within 1 km^2^ in a mixed hardwood forest on scales from centimeters to tens of meters, we detect persistence over 3 years of native genotypes, identified by SNPs genome-wide, of the wild yeast, *Saccharomyces paradoxus* around *Quercus rubra* and *Q. alba*. Yeasts were recovered by enrichment in ethanol-containing medium, which measures only presence or absence, not abundance. Additional transplantation experiments employed strains marked with spontaneous defects in the *URA3* gene, which also confer resistance to 5-Fluoroorotic acid (5FOA). Plating soil suspensions from transplant sites on 5FOA medium permitted one-step quantification of yeast colony-forming units, with no interference from other unmarked yeasts or microorganisms. After an initial steep decrease in abundance, the yeast densities fluctuated over time, increasing in association with rainfall and decreasing in association with drought. After 18 months, the transplanted yeasts remain in place on the nine sites. In vitro transplantation experiments into non-sterile soil in petri dishes showed similar patterns of persistence and response to moisture and drought. To determine whether *S. cerevisiae*, not previously recovered from soils regionally, can persist in our cold-climate sites, we transplanted marked *S. cerevisiae* alone and in mixture with S. *paradoxus* in fall, 2017. Five months on, *S. cerevisiae* persist to the same extent as *S. paradoxus*.

## IMPORTANCE

Saccharomyces yeasts are intensively studied in biological research and in their domesticated roles of brewing and baking. And yet, remarkably little is known about their mode of life in forest soils. We report here that resident genotypes of the yeast *S. paradoxus* are persistent of a time scale of years in their micro-habitats in forest soils. We also show that resident genotypes can be replaced by transplanted yeast genotypes. The high inoculum levels in experimental transplantations rapidly decreased over time, but the transplanted genotypes persisted at low abundance. We conclude that, in forest soils, Saccharomyces yeasts exist at very low abundance and that dispersal events are rare.

In their review of the ecology and evolution of non-domesticated *Saccharomyces* species, Boynton and Greig (1) encouraged further investigation in nature to place them in ecological context with reference to the scientific model, *S. cerevisiae*, long-domesticated for food and beverage fermentation. Our focus here was on *S. paradoxus*, the most tractable comparator of wild to domesticated *Saccharomyces. S. paradoxus* is associated with, but not limited to, oak leaf litter in forest soils (2, 3). In this yeast, progress has been made on understanding the relative importance of dispersal, genetic drift, and local adaptation in populations (4), and the effects of substrate utilization on metabolism and fitness (5, 6). Saccharomyces yeasts are present in soil and plant material, especially bark of deciduous trees where they utilize exudates such as sap. Filteau et al. (6) have elucidated the genetic basis of a fitness determinant in *S. paradoxus*: degradation of allantoate, the major nitrogen source in maple sap. However, there has been little information on absolute abundance of these yeasts in their natural habitat. Most information on the occurrence of wild yeasts in soil has come through enrichment culture in ethanol-containing medium, which cannot be used to estimate abundance. A major goal of this study was to measure the abundance over time of yeasts transplanted to their natural substrates at high initial densities.

Dispersal of *Saccharomyces* species between substrates is poorly understood (1), but the recent work of Boynton et al. (4) suggests that lack of dispersal and prominence of genetic drift are more important in shaping local populations of *S. paradoxus* than fitness differences among genotypes. *S. paradoxus* may be dispersed by rainwater and by insects such as *Drosophila* (7, 8), from which it has been isolated. Despite these potential dispersal mechanisms, there is genetic differentiation among *S. paradoxus* populations that is roughly proportional to distance on a scale from centimeters to thousands of kilometers (9). Despite progress in describing population-genetic structure and in documenting potential dispersal mechanisms, the extent to which wild yeasts actually grow and persist in stasis in their habitats in soil is unknown.

In this study, we first followed naturally occurring genotypes of *S. paradoxus* over a three-year period on a fine geographical scale of marked sites in a natural woodland. This extended the time frame covered by our earlier study (10) by two years. This part of the study included only enrichment culturing and therefore registered only presence or absence, not abundance of yeasts. We then initiated transplantation experiments with yeast strains marked with spontaneous mutations. The transplanted yeasts in our study allowed quantification of colony-forming units per unit of soil and of change over time from an initially high level of inoculum. We found that abundance fluctuated, with an overall downward trend over time and that rainfall events were associated with a temporary increase in abundance. Even when abundance counts approached zero in some transplant sites, enrichment culture invariably recovered the transplanted genotype. Although our experiments focused on *S. paradoxus*, we show that marked strains of *S. cerevisiae*, although not among the residents found in this locality, persist in proximity to transplanted *S. paradoxus*.

## RESULTS & DISCUSSION

In 2014 and 2015, yeasts were collected by enrichment culture in ethanol medium from 72 sites around the bases of three oak trees and genotyped for SNP sites across their genomes. A remarkable pattern of persistence of genotypes in their sites of origin was observed over this one-year time span. In the present study, we sampled these 72 sites again in 2016 and a limited selection of eight sites again in 2017 (Table 1), extending the sampling times of 2014 and 2015 reported by Xia et al. (10) by two years.

**Table 1.**
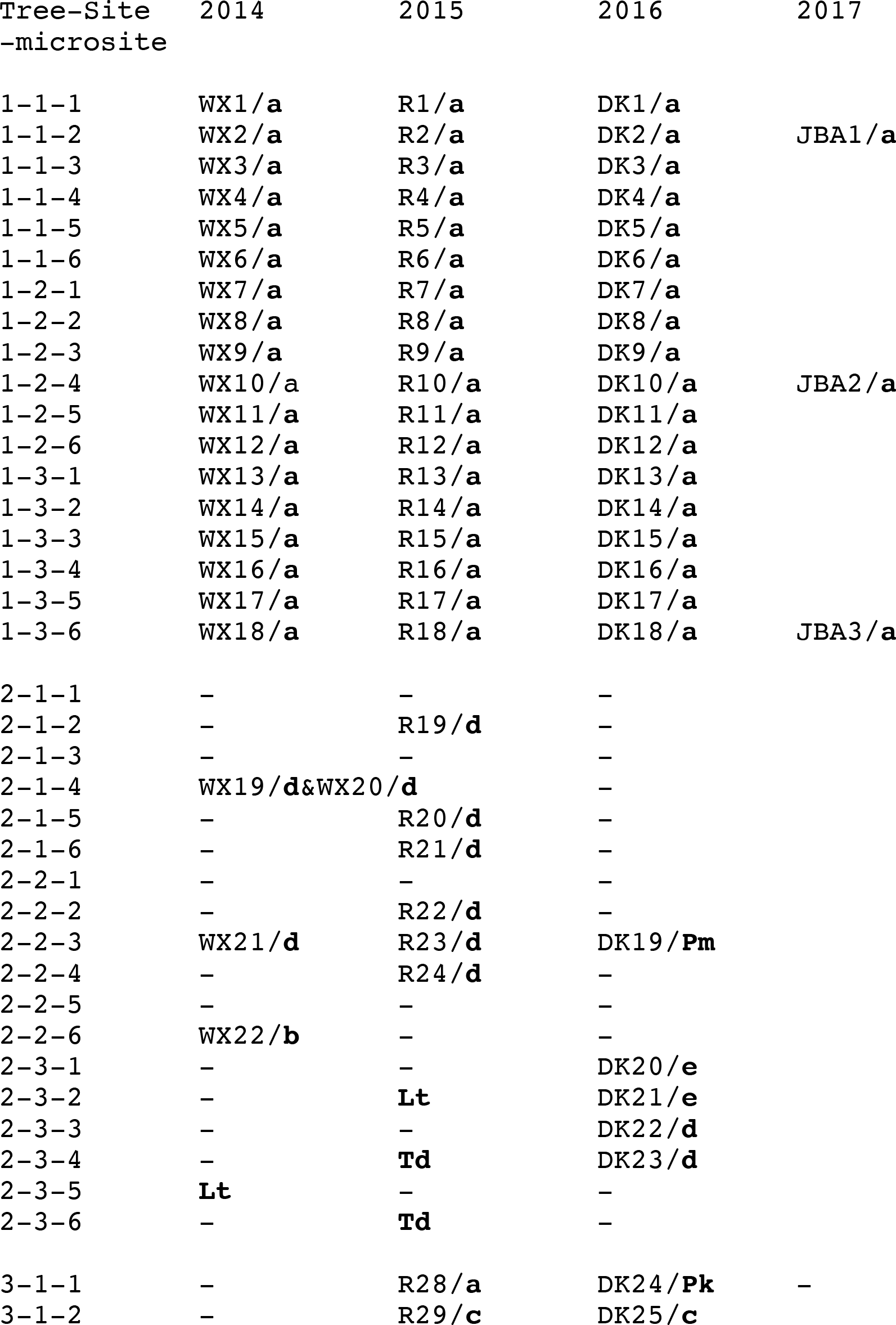
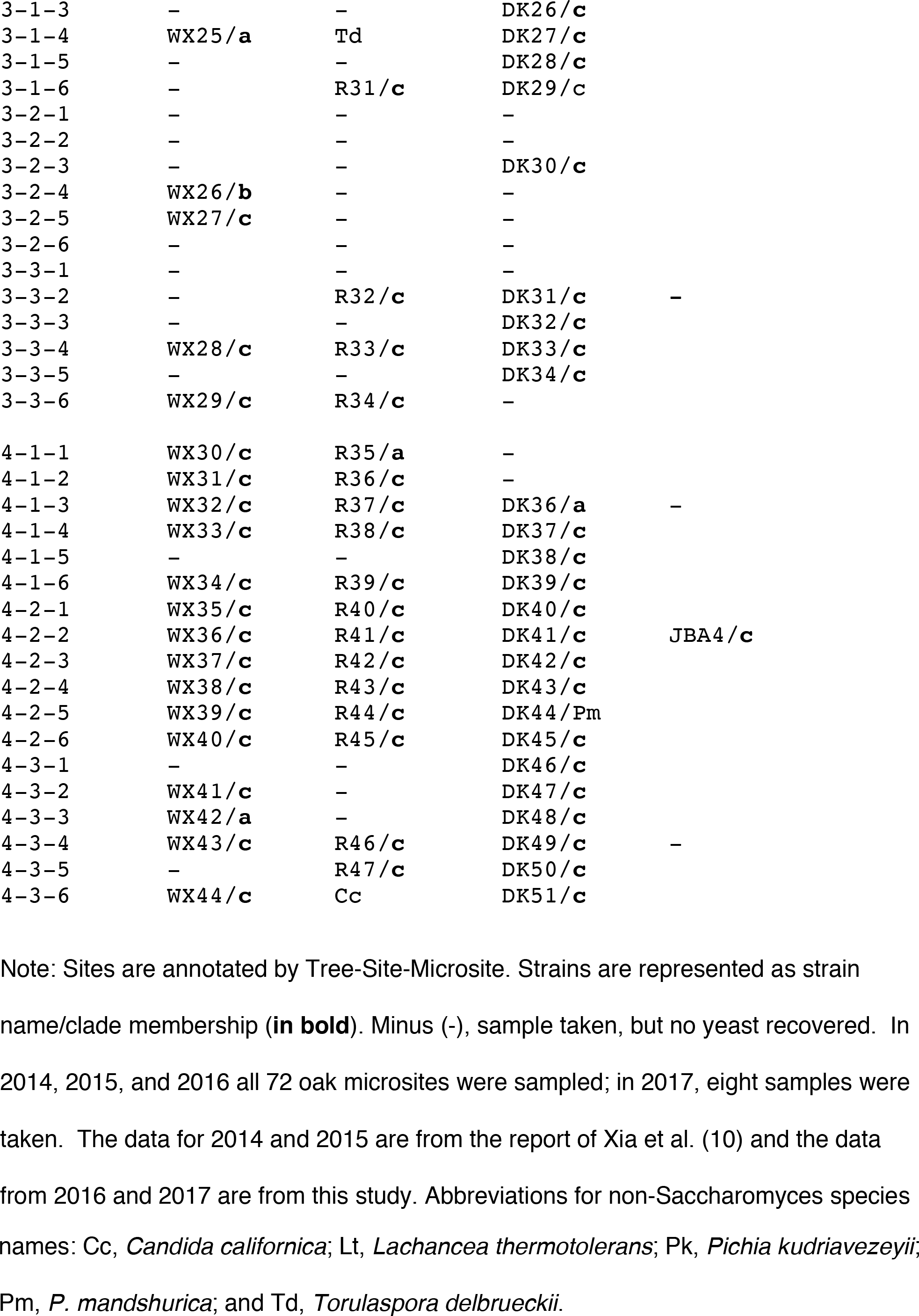
Identity of yeast strains recovered by enrichment culture from sites near the bases of four oak trees.

At Oak 1, all sites were occupied in all years by a single *S. paradoxus* genotype, clade a. Here, the rate of recovery of *S. paradoxus* was remarkable in its completeness across all sites. Oak 2 showed an entirely different pattern of occupancy from that of Oak 1. Overall, less than half the sites at Oak 2 returned yeasts, but those that did included a broad diversity of genotypes, including: *S. paradoxus* clade d, a hybrid lineage between two divergent lineages; an isolate of clade b, also a hybrid, but between more recently diverged lineages (clades a and c); clade e, a genotype of European origin, and three other yeast species, *Lachancea thermotolerans, Torulaspora delbrueckii*, and *Pichia mandshurica*. Oak 2 is also notable for the complete absence of clades a and c among the recovered yeast samples.

Oak 3 and Oak 4 presented partial occupancy of sites, a predominance of clade c among the positive sites, and minority of clades a, b. There were three additional yeast species: *Pichia kudriavezeyii, P. mandshurica*, and *Candida californica*, a species reported to be consistently vectored by *Drosophila melanogaster* (11). The final year of limited sampling yielded one isolate of clade c. Overall, the pattern of persistence is extended at least through 2016, and is consistent with persistence in the limited sample into 2017.

The previous re-sampling experiments examined yeasts in natural sites around the bases of oak trees by enrichment in ethanol containing medium; this culture method is not quantitative and only registers presence or absence, with some unknown threshold for detecting presence. In studying natural populations of *S. paradoxus*, there has been a clear need for a method to quantify transplanted yeasts in situ. A recently reported method for quantifying yeasts in their natural habitats (4) used digital droplet PCR, which sensitively measures the ratios of the abundance of transplanted genotypes, but not their absolute abundance. In this study, we used strains marked with spontaneous mutations in the *URA3* gene, which conferred a Ura-, 5FOA-resistant phenotype. With this method, absolute abundance, colony forming units per g of soil, could be measured in one step by dilution plating on agar medium containing 5FOA, with no interference by growth of other soil microorganisms.

We transplanted yeast populations of high density into nine sites around the bases of three additional oak trees (Oak 6, Oak 7, and Oak 8) in order to measure their abundance over time in the natural habitat. Figure 1 shows the change in the composite abundance of all transplant populations over time. After a steep decrease in abundance over the first two weeks after transplantation, the levels fluctuated over time with seven of nine sites still registering counts one full year after transplantation and two of nine sites returned no CFUs. However, enrichment cultures at the end of the experiment, including those two sites registering no CFUs in the final plating on 5FOA medium, were all positive for the transplanted strains. The original resident yeasts on these sites sampled before transplantation (Table S1) would not be expected to appear on these 5FOA plates; whether or not the original resident strains remain on the transplant sites at low frequency is not known.

**Figure 1.**
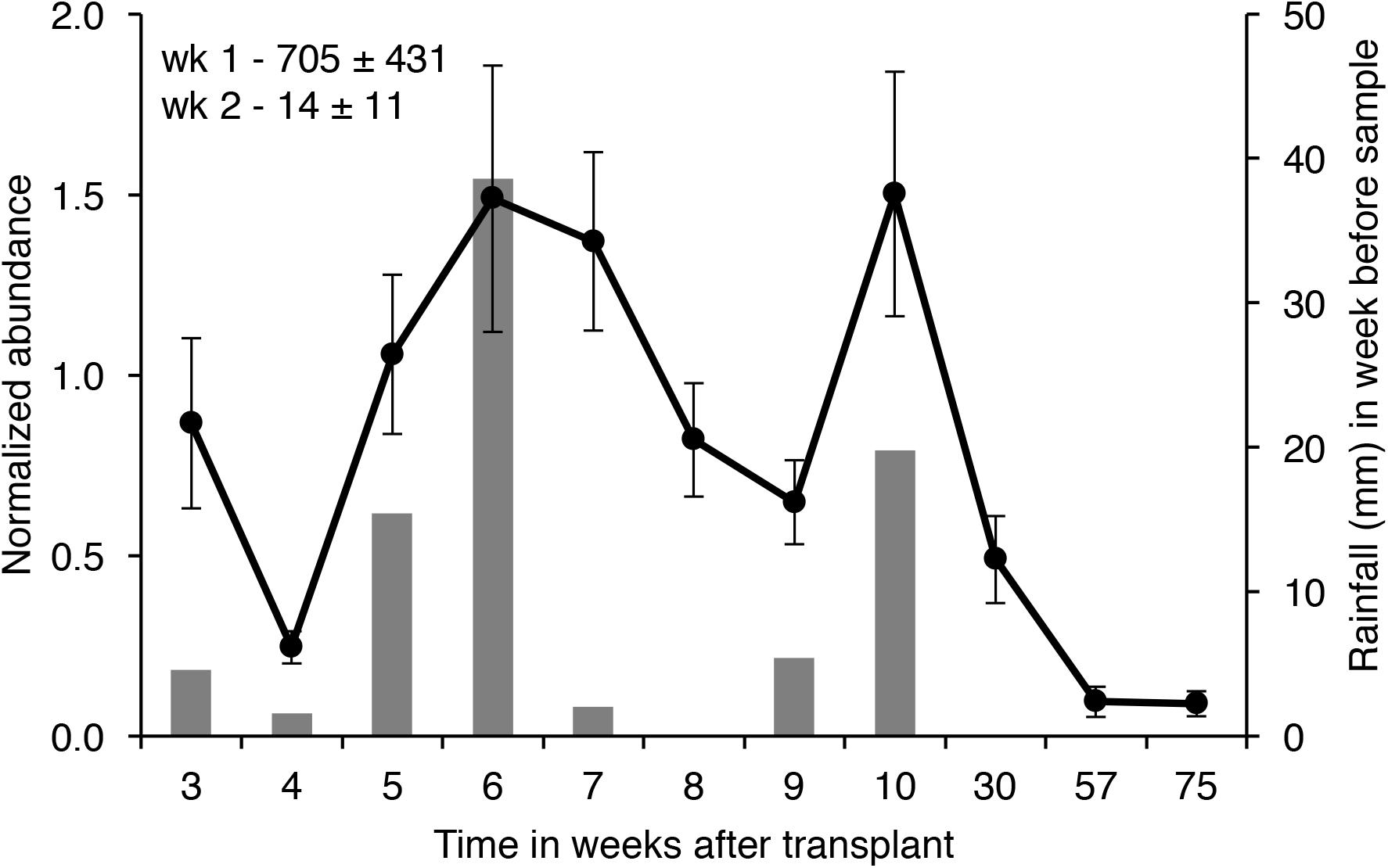
Outdoor transplantation of yeast strains and abundance over time. Three strains were each transplanted to three sites around the bases of three oak trees (three strains per tree). The abundance values for each site were normalized against the average counts of colony forming units (CFU) per gram of soil between weeks 3 and 8 plus or minus standard error. The average value for normalization (over the nine sites) was 373,440 CFU / g soil ± 76,254 (SE). In the initial sampling, the colonies were too numerous to count on the dilution plates. The average relative abundance values for all nine sites, plus or minus standard error for weeks 1 and 2, appear in the upper left; these values are off the scale of this graph. The bars represent rainfall amount in the week prior to sampling.

In addition to abundance, Figure 1 also indicates rainfall over time. For the first and larger of the two conspicuous rainfall events, between weeks four and six, all (nine of nine) populations increased in abundance (under the null hypothesis that increase in CFUs, or not, is a random behavior, the probability of the same directionality of response in all nine sites: P ≤ 1/ 2^9^, or 0.002). In the second event, between weeks 8 and 10, eight of the nine populations increased in abundance. At this stage, we interpret the apparent increase in population abundance with rainfall as a possible association, which is addressed below in a laboratory experiment. This transplantation experiment was not expected to distinguish whether the increase was due to reproduction of cells or to separation of aggregated cells because of moisture (driving up CFUs on the detection plates), or to some other mechanism.

The outdoor transplants introduced yeast populations to a spot about 5 cm in diameter. At week 39, we recovered samples along six transects extending outward from three transplant sites (Figure 2). Here the abundance values clearly show that the plume of transplanted yeast cells had spread laterally. How this happened cannot be distinguished in this experiment. The yeast cells may have been washed outward from their central location with rainfall. This lateral spread may in part explain rapid initial decrease in abundance in the central location after transplantation. Actual reduction in viable cells may also contribute to the reduction in observed CFUs.

**Figure 2.**
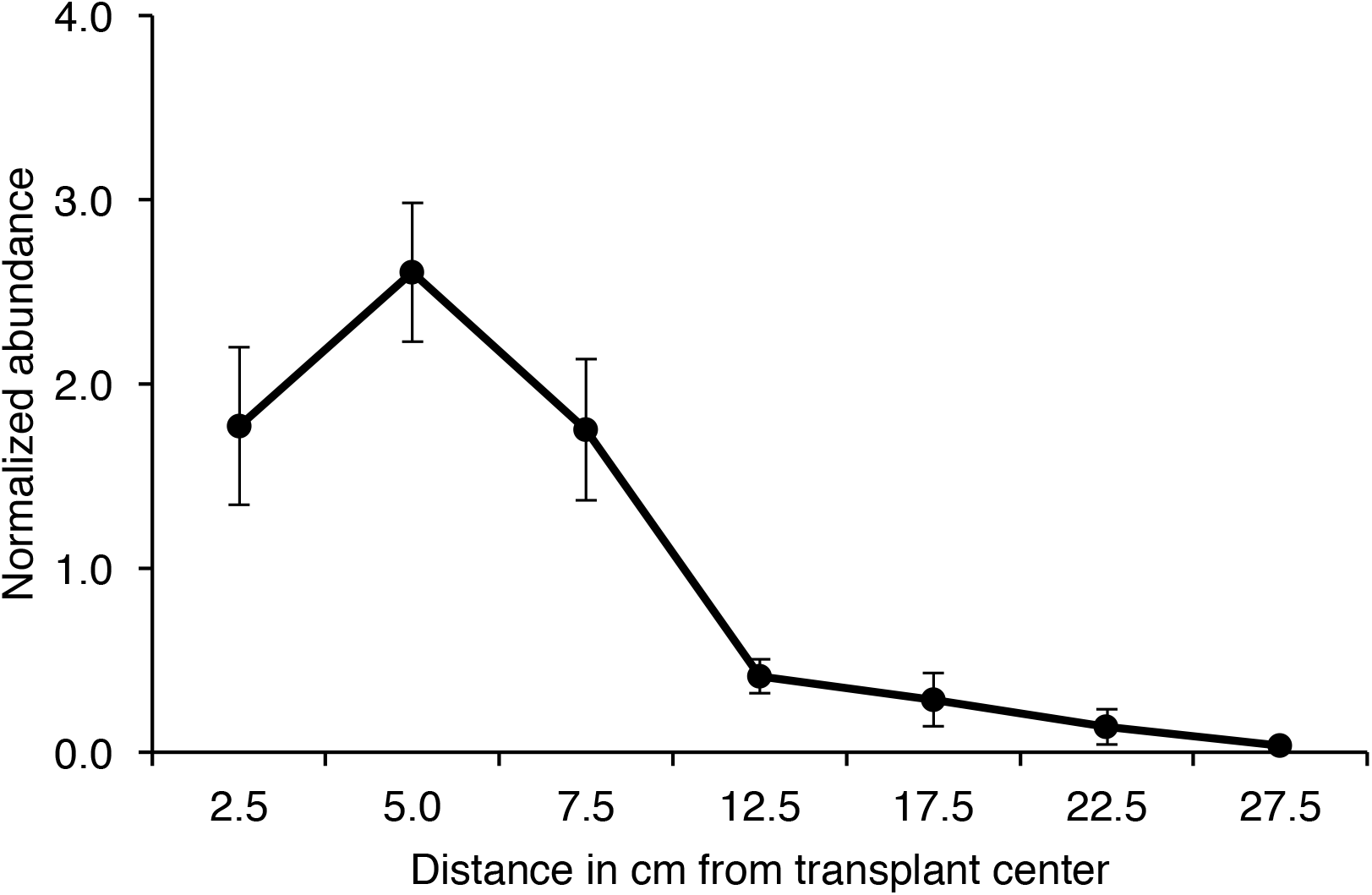
Outdoor transplantation and relative abundance with distance in cm from the transplant site. On three of the nine sites for outdoor transplantation (each at a different tree and each a different strain), six diameters were measured at distance intervals (two intervals per site) at week 39 after transplantation. The values for each site were normalized against CFU / g soil for each of the six diameters plus or minus standard error. The average value for normalization (over the six diameters) was 58,449 CFU /g soil ± 14,644 (SE).

The next experiment was devised to reproduce the outdoor transplant experiment in vitro, with petri dishes containing a limited volume of non-sterile soil. The pattern of abundance over time in the indoor experiment paralleled the outdoor experiment. Yeast abundance initially decreased sharply and then fluctuated over time, gradually approaching low levels by week 41 (Figure 3). A final enrichment culturing at week 48, followed by testing on 5FOA medium, revealed that all nine plates harbored viable representatives of the originally transplanted strains. Under these conditions, the decline in CFUs could not have been due to spreading of the yeast populations as occurred in the outdoor transplantation sites. The decline of CFUs must have been due to a decrease in the proportion of viable cells or in their propensity to form aggregates (in which multiple viable cells that were aggregated would plate as one CFU).

**Figure 3.**
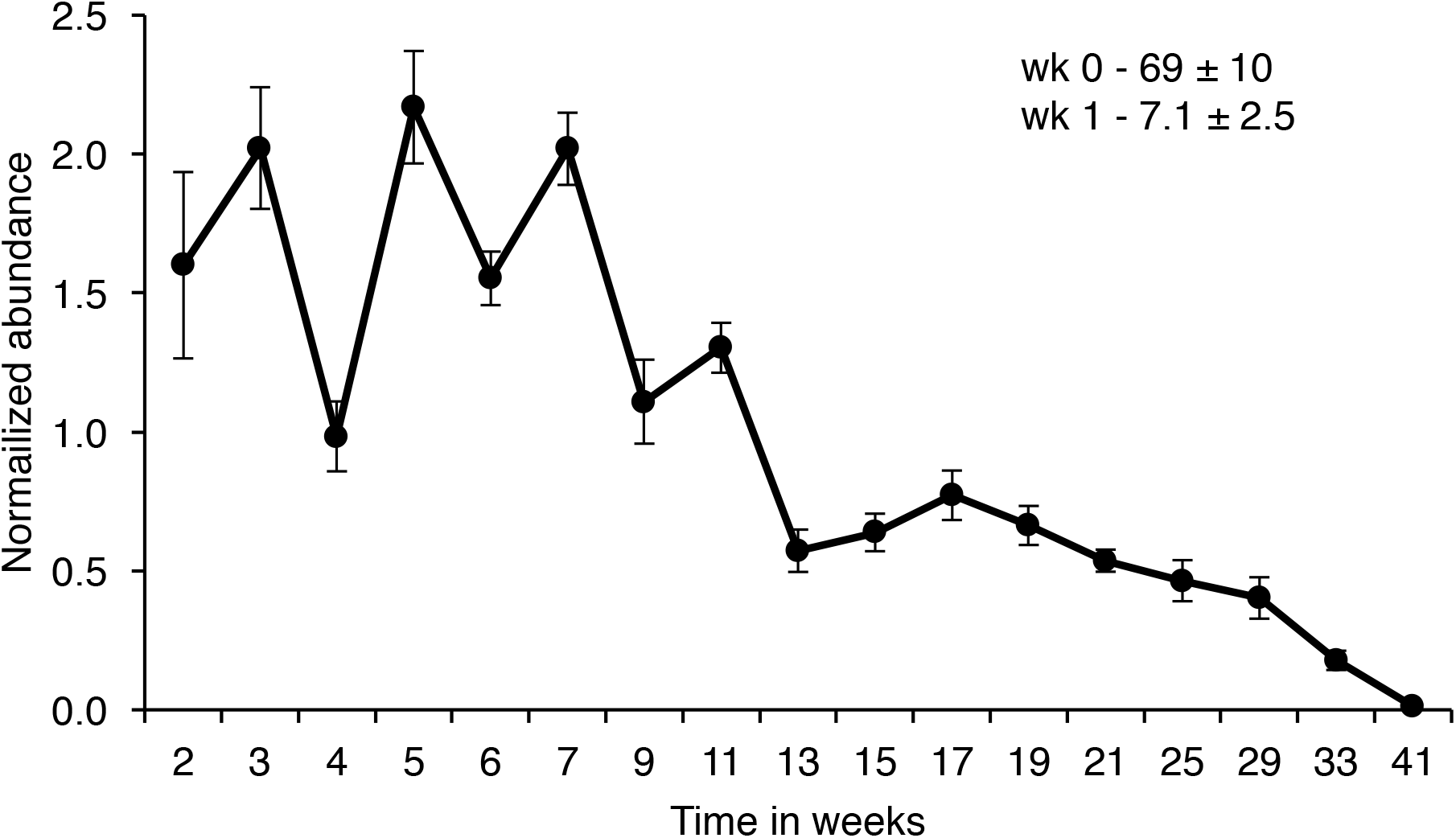
Indoor transplantation of yeast strains and abundance over time. Three strains were transplanted to each of three petri dishes containing 50 g of forest soil (nine plates total). The values for each site were normalized against CFU / g soil between weeks 2 and 41 plus or minus standard error. The average value for normalization was 612,118 CFU / g soil ± 74,280 (SE). The values for the initial sampling and week 1 after transplantation appear in the upper right.

After the indoor transplantation experiment was completed, the soil plates were re-purposed for another experiment to test for the effect of moisture after extended drought. The plates were allowed to dry out at room temperature for three months. At this point, soil samples were taken for dilution plating on 5FOA medium and for enrichment culture. No CFUs were registered for this initial sampling; the transplanted yeasts had become rare. Nonetheless, the enrichment cultures were all positive for the transplanted yeasts (which had the 5FOA phenotype). Water was then added to the plates so that the soil became saturated, but without free water. Water was added to the plates twice per week to keep the soil saturated. Samples were taken at week 1 and week 2 two after the initial sampling. All plates registered CFUs and, for all replicates, the CFUs per g soil increased from initial sampling (0.0 ± 0.0 SE) to week 1 (1367 ± 519 SE), and to week 2 (2156 ± 764 SE). The effect in this in-vitro experiment was remarkably consistent with response to rainfall after a period of dryness in the outdoor transplantation experiment (null hypothesis, increase in CFUs, or not, is random; probability of the same directionality of response in all nine plates: P ≤1/ 2^9^, or 0.002).

In a final experiment, we addressed the question of whether *S. cerevisiae* would persist in outdoor transplant sites, as did *S. paradoxus*. The rationale for this experiment is that *S. cerevisiae* is not found in our study site or in northern latitudes in woodlands (12, 13). Furthermore, genotypes of *S. cerevisiae* do not appear to spread from vineyards to forests (14). Here, we transplanted a marked genotype of *S. cerevisiae* in proximity to *S. paradoxus*. Both yeast species and the mix declined in abundance over the first six weeks to very low CFU counts at the end of the winter (week 18, Figure 4). The ratio of the two yeast species fluctuated around 1:1. At the end of the experiment CFUs were low (Table S2); in the mixed population, in total, there were six CFUs of *S. cerevisiae* and six of *S. paradoxus*. In enrichment culture at week 19, all nine sites recovered the original genotype transplanted. At this stage, *S. cerevisiae* appears to persist about as well as *S. paradoxus* (5) in the study locality, where they have not been detected previously.

**Figure 4.**
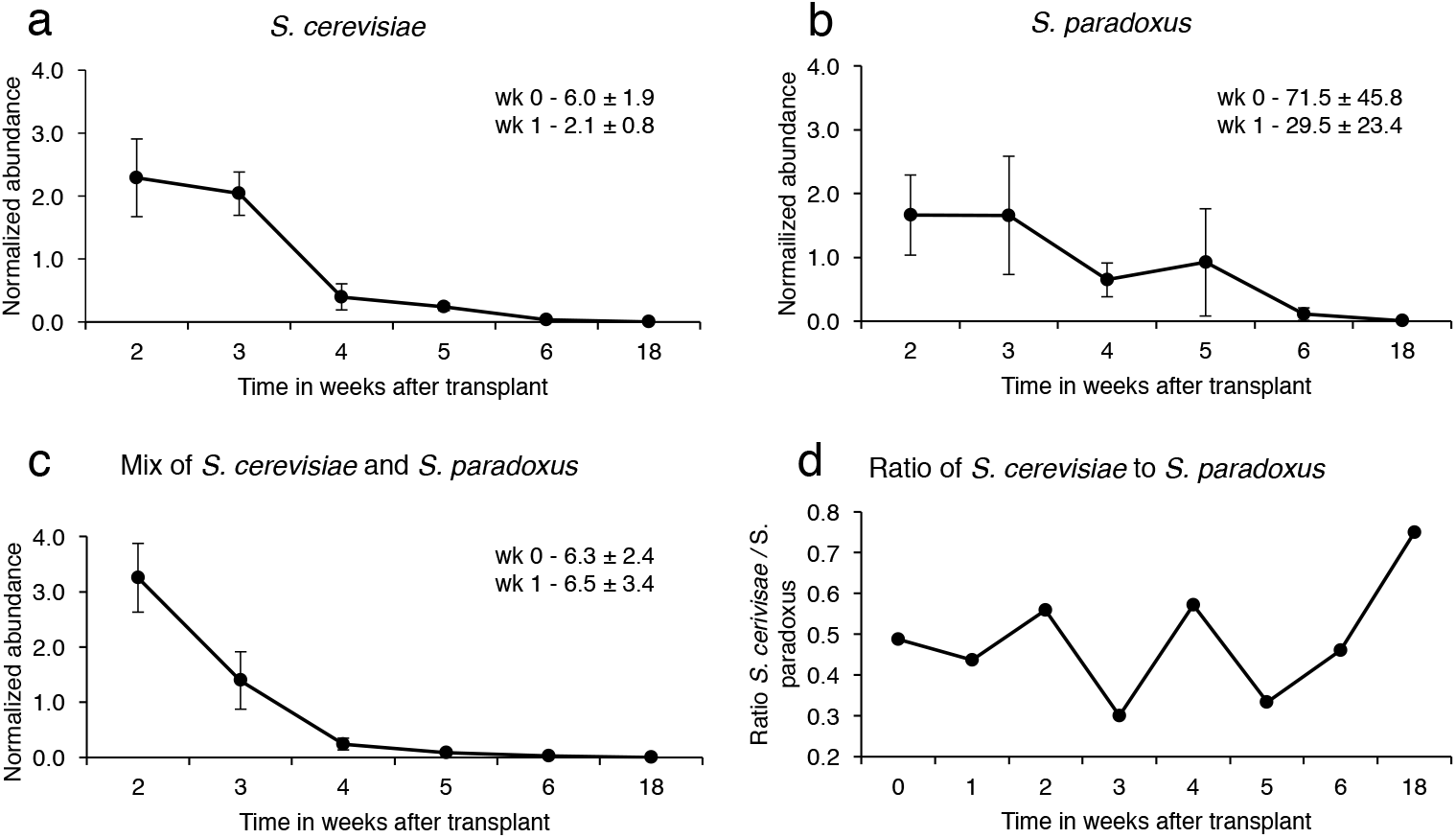
Outdoor transplantation of *S. cerevisiae* and *S. paradoxus* and relative abundance over time. Inocula containing (**a**) *S. cerevisiae* only, (**b**) *S. paradoxus* only, and (**c**) a mix of *S. cerevisiae* and *S. paradoxus*, were transplanted to three sites at the bases of three oak trees; each tree had one site of each type of inoculum, (**d**) the ratio between *S. cerevisiae* and *S. paradoxus* for the mixed population; note that counts on 5FOA test plates were low for the 18-week time point, total of 12 CFUs on nine plates. The values for each site were normalized against CFU /g soil between weeks 3 and 10 plus or minus standard error. The average values for normalization were: *S. paradoxus*, 738;,722 ± 632,755 (SE); *S. cerevisiae* 922,523 ± 592,478, and the mix of the two yeasts 1,287,051 ± 315,193 (SE). The average relative abundance values (all nine sites) for weeks 1 and 2, which were off scale on the graph, appear in the upper right.

There are two main contributions from this study. The first contribution is technical. Although our focus here was on persistence of yeast strains in their habitats, our transplantation experiments demonstrate the feasibility of long-term measurements of fitness effects in mixtures of strains in the field and in the laboratory. An advantage of transplantation to indoor microcosms with non-sterile soil is that the markers used to distinguish strains need not be limited to spontaneous mutations as was the case here for outdoor transplantation experiments. Targeted comparisons might involve specific gene deletions or modifications that would not be appropriate for outdoor transplantation.

The second contribution is biological. Our observations that resident yeast populations persist over time with infrequent dispersal is entirely consistent with the scenario proposed by Boynton et al. (4). Our transplantation experiments are also consistent with a low level of dispersal, contributing a new dimension to our understanding of the carrying capacity of soil habitats for yeast populations. The initial high levels of inoculum in the transplantations were not sustainable. The CFUs per g of soil decreased precipitously in the first two weeks after transplantation and then more slowly, finally registering few or no CFUs on the 5FOA selective medium. And yet, enrichment culturing indicated that the original, transplanted genotypes were still on their sites in a viable state. Although the initial high levels of inoculum were not maintained, the transplanted genotypes persist and the transplants replace the naturally occurring yeast residents on those sites. Only by sampling the transplant sites in subsequent years can the limits of persistence be measured. Overall, our results are consistent with the idea that Saccharomyces yeasts exist at extremely low levels in the soil, with the possibility for limited population growth when temperature and moisture conditions in the soil permit. Against this prevalent pattern of stasis, the sudden availability of higher nutrient conditions, for example from sap flows, could lead to substantial growth in yeast populations with subsequent decrease in population density over time; our transplantation experiments were designed to simulate these high-growth events. We speculate that such chance amplifications in yeast populations would have more effect on long term presence than fitness differences among genotypes, especially where those fitness differences only relate to slight differences in growth rate measured under rich nutrient conditions in laboratory culture.

## MATERIALS AND METHODS

### Enrichment recovery of naturally occurring yeasts

Our initial goal was to identify naturally occurring genotypes of *S. paradoxus* and to track their distributions over time. The sampling area included the same sites studied via enrichment culture by Xia et al. (10) in 2014 and 2015 and this study extended the earlier sampling into 2016 and 2017 (Table 1). The sample encompassed four oak trees, with three sites around the base of each tree separated by buttress roots, with six microsites in each site (Fig. 1, See Xia et al., 10). Collection of soil samples, enrichment culture with 8% ethanol, DNA isolation, Illumina sequencing, read alignment against a standard genome, and variant discovery were exactly as described by Xia et al. (10). Each newly isolated strain was identified to clades a – e, which were described earlier by Xia et al. (10)

### Marked strains for transplantation experiments

Transplantation experiments were done with spontaneously arising *ura3* mutants. We found these mutants in *S. paradoxus* by allowing a diploid culture to sporulate and then plating large numbers of spores in tetrads on 5FOA medium. The sporulation step was done to facilitate homozygosity for any newly arising mutants. The resistant mutants appearing on 5FOA medium had a Ura-phenotype (15). Three *ura3* mutants of *S. paradoxus* were used: one frameshift (Sce5003, c.260_261insC), one nonsense (Sce5006 c.553C>T, GLU to stop), and one missense mutation (Sce5010, c. 801G>A, GLY to ASP).

For a marked version of *S. cerevisiae*, we crossed a wild type strain (Sce13) with another strain carrying multiple auxotrophies (Sce695). From the offspring of this cross, we identified haploid strains of complementary mating type that were 5FOA resistant (and Ura-) and His-. These two haploids were then mated to form a diploid that was homozygous for *ura3* and *his3*. The *his3* mutation was used to distinguish *S. cerevisiae* from *S. paradoxus* (His+) in mixed transplant sites. None of the transplanted strains of either *S. paradoxus* or *S. cerevisiae* contained any genetically engineered, or recombinant DNA, constructs.

### Outdoor transplantation of *S. paradoxus*

Oak 6, Oak 7, and Oak 8, were selected for transplantation. At each tree, there were three transplant sites separated by buttress roots (total of nine sites). The three sites at each tree were inoculated with a different one of the three *ura3*- marked strains of *S. paradoxus* (23 Sept., 2016). Immediately before transplantation, any yeast residents were identified by enrichment culture as described (Table S1).

At each site, we applied 100 ml of water with a suspension of 10^10^ cells through the perforated cap of a large saltshaker. A spot on the soil surface of about 5 cm in diameter was sprinkled with the cell suspension, allowing sufficient time for the fluid to soak in without running across the surface. At the time of sampling, ca. 0.5 g of soil was scraped into a 15 ml plastic tube, weighed, and suspended in either 10 ml of water or, at the end of the experiment, in 10 ml of enrichment medium (6). After vigorous agitation, 100 *μ*l of the suspension, or a dilution thereof, was spread onto 9 cm petri dishes with 5FOA medium (15). After three days of incubation at 30 C, colonies were counted and the number of CFUs per g of soil calculated. Raw CFU data are given in Table S2. At the end of the experiment, and after enrichment culturing, colonies were tested for 5FOA resistance and the URA-phenotype. Sites without transplantation invariably registered no 5FOA resistant cultures.

### Indoor transplantation of *S. paradoxus*

These experiments were constructed to mimic the outdoor experiments, except that the available volume was limited and the inoculum was initially mixed into the non-sterile soil, which had been collected near the bases of several oak trees, pooled, and sieved to a fine particle size. Deep petri dishes (2.5 × 9 cm) containing ca. 50 g soil mix were inoculated with 10 ml of water containing a suspension of 10^9^ cells (12 Jan., 2017). Each soil plate was weighed initially and then weekly thereafter; the reduction in weight over the preceding week was compensated by the addition of sterile water. After this experiment was completed, the soil plates were re-purposed to test the effect of moisture addition after a period of extended dryness as described below. The plates were allowed to dry out at RT for three months and CFUs were measured before and after the addition water to the plates.

### Outdoor transplantation of *S. cerevisiae*

We selected three new trees (Oak 9, Oak 10, and Oak 11) with three sites around the base of each tree separated from one another by buttress roots. At each tree, *S. cerevisiae* was transplanted to one site, *S. paradoxus* to another site, and an equal mix of the two species to a third site. Transplantation yeast cells was done exactly as in the outdoor transplantation experiment. Approximately 10^10^ cells suspended in water were applied to each site (26 October, 2017).

### Data availability

A comprehensive variant (.vcf) file for all *S. paradoxus* strains in Table 1 is available in the Dryad Digital Repository (accession number pending acceptance). Alignments of Illumina reads of the 2014 and 2015 collections with a reference *S. paradoxus* genome (.bam files) are accessible through NCBI’s SRA (http://www.ncbi.nlm.nih.gov/sra) as accession PRJNA324830 (see ref. no. 10).

## SUPPLEMENTAL MATERIAL

**Table S1**. Summary of resident yeast species on the sites for outdoor transplantation.

**Table S2**. Raw data (CFUs / g soil) for Figures 1–4.

## ACKNOWLEDGEMENTS

This study was supported by Individual Discovery Grants from the Natural Sciences and Engineering Research Council of Canada (NSERC) to JBA and LMK.

